# Responsive robotic prey reveal how predators adapt to predictability in escape tactics

**DOI:** 10.1101/2021.05.12.443728

**Authors:** Andrew W. Szopa-Comley, Christos C. Ioannou

**Affiliations:** School of Biological Sciences, Life Sciences Building, 24 Tyndall Avenue, University of Bristol, Bristol, BS8 1TQ, U.K.

## Abstract

To increase their chances of survival, prey often respond to predators by being unpredictable when escaping, but the response of predators to such tactics is unknown. We programmed interactive robot-controlled prey to flee from an approaching blue acara predator (*Andinoacara pulcher*), allowing us to manipulate the predictability of the prey’s initial escape direction. When repeatedly exposed to predictable prey, the predators adjusted their behaviour before the prey even began to escape: prey programmed to escape directly away were approached more rapidly than prey escaping at an acute angle. These faster approach speeds compensated for a longer time needed to capture such prey during the subsequent pursuit phase, and predators showed greater acceleration when pursuing unpredictable prey. Collectively, these behaviours resulted in the prey’s predictability having no net effect on the time to capture prey. Rather than minimising capture times, predators adjust their behaviour to maintain an adequate level of performance.

## Introduction

Rapid evasive responses are a vital tool prey use to minimise capture by predators (1, 2). Despite their ubiquity, it can be challenging to demonstrate the benefit of escape strategies because integrating realistic predation and manipulation of prey behaviour that experimentally controls for confounding effects is difficult. Studying the behaviour of real predators is crucial when attempting to demonstrate the adaptive value of prey adaptations, especially when these are dependent on features of predator cognition (3-5). This applies particularly to unpredictable escape behaviour by prey, which is thought to enhance prey survival by compromising the ability of predators to anticipate the movement of their target (6). Although unpredictable escape tactics are widespread taxonomically (7, 8), we know little about how real predators respond to unpredictability in prey escape strategies, and whether this prevents predators from adjusting their behaviour over multiple interactions with prey (9, 10).

Controlled experiments in which human ‘predators’ target continuously moving virtual prey have demonstrated that abrupt and unpredictable changes in direction enhance prey survival by reducing the accuracy of prey targeting (11, 12). However, it is unknown whether the survival advantage conferred by unpredictable motion also applies against non-human predators, or to escape responses of prey which are initially stationary rather than in continuous movement. This is common in nature, as numerous prey taxa freeze once they have detected a potential threat or remain motionless to avoid detection by predators, before eventually fleeing only once a predator gets too close (1, 13-15).

One way for stationary prey to be unpredictable over successive interactions with predators is to vary the initial escape angle from one encounter to the next (16). Although theoretical models predict that prey should select a single optimal escape trajectory which maximises the distance from an approaching predator (17, 18), predators might be able to learn to anticipate the movements of prey which repeatedly escape at a fixed angle relative to their approach (19). Initial prey escape angles are often surprisingly variable (20-25), and while it has been suggested that this is due to biomechanical or sensory constraints on the angle of escape, the variability in escape angles might also represent an anti-predator strategy aimed at generating unpredictability (26). Varying the initial angle of escape could prevent predators from learning to anticipate the directional heading of their target over multiple attacks (16, 26, 27), but previous studies on prey escape behaviour have not tested how real predators respond to unpredictable tactics.

Many predator-prey interactions are typified by feedback between the attacker and the target (28), making it difficult using a purely observational approach to disentangle the effects of prey defences on predators from the impact of predator behaviour on prey. This lack of experimental control can be avoided by presenting real predators with standardised virtual prey, whose movements and behaviour can be precisely controlled and experimentally manipulated (29-33). However, previous experiments with virtual prey have used unresponsive prey items which do not react to the predators, and do not allow the predator to capture prey and be rewarded. These limitations can be overcome by using interactive robotic prey (34), making it possible to investigate predator responses to prey escape behaviour.

To study the effect of unpredictability in prey escape on predators, we develop a novel experimental system (**Fig. 1a**) in which artificial robot-controlled prey items were programmed to flee from blue acara cichlid (*Andinoacara pulcher*) predators once the predator had approached within a threshold distance. Blue acaras are opportunistic predators with a broad diet, but often actively pursue highly evasive prey such as Trinidadian guppies (35, 36). After an initial period in which groups of blue acaras were trained to attack the prey (the training period), individual predators were assigned to one of two experimental treatments and repeatedly presented with prey over twenty successive experimental trials (the test period). Prey in the two treatments differed in the consistency of their escape behaviour from trial-to-trial: in the predictable treatment, prey escaped in the same direction relative to the predator’s approach from one trial to the next, whereas in the unpredictable treatment, prey were programmed to flee in random directions over successive trials (**Fig. 1b-c**). To successfully capture prey, pursuit predators must respond to changes in prey direction which occur at the start of a chase (37-39). Across trials with predictable prey, predators had the opportunity to gain reliable information about the prey’s likely escape direction, but in the unpredictable treatment, the escape angle of the prey in previous trials was not a reliable indicator of its escape direction in future encounters.

**Figure 1:**
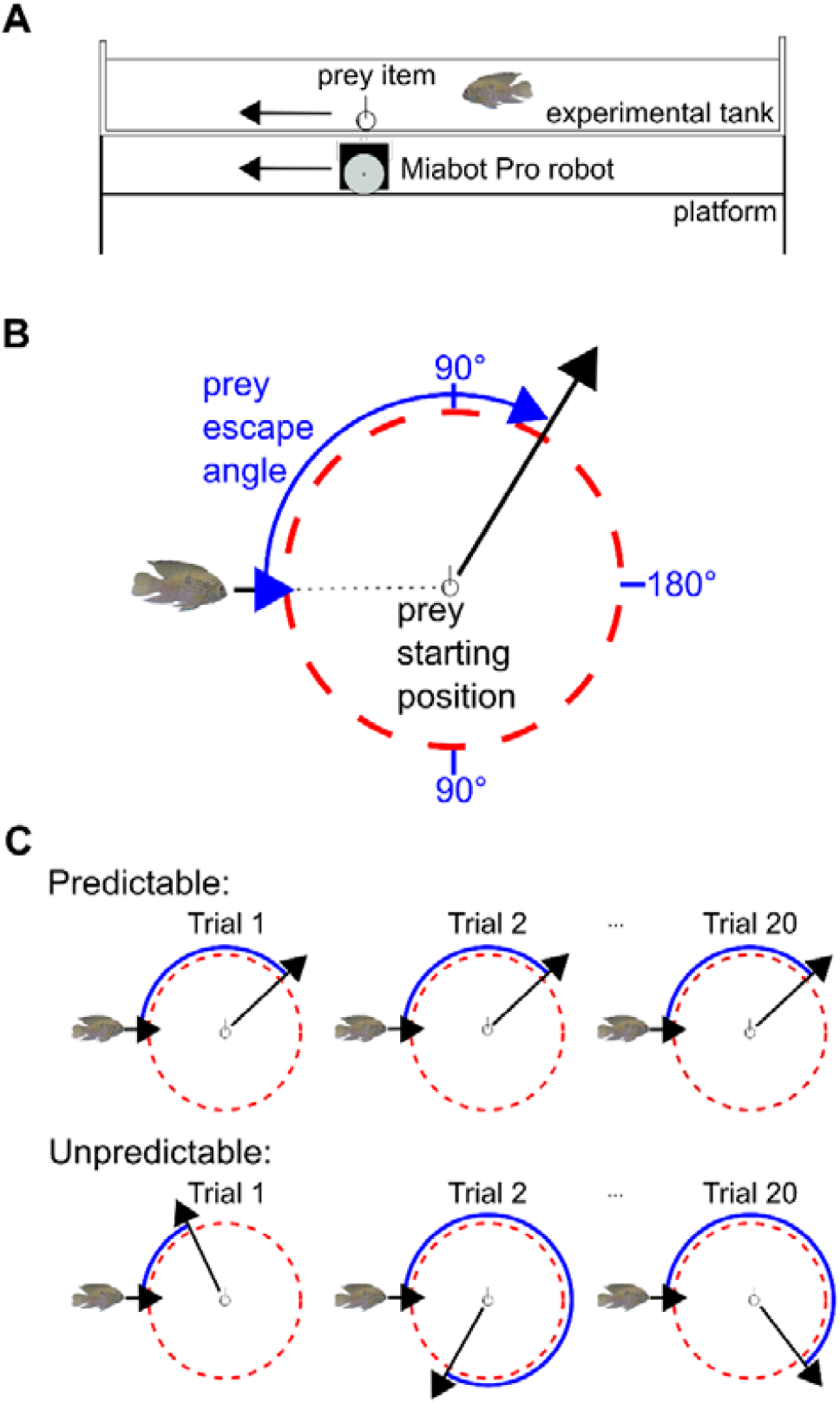
The robotic prey system. (a) Diagram (not to scale) showing a side-view of the experimental tank, with the Bluetooth-controlled robot situated on a platform underneath the experimental tank. The robot controls the movements of the artificial prey item via magnets. (b) The prey’s escape angle was defined relative to the predator’s approach direction. (c) In the predictable treatment, individual predators were exposed to prey which escaped at the same angle over successive trials (the escape angle varied between individual predators). In the unpredictable treatment, the prey escape angle varied randomly from trial to trial.

## Results

### Performance of the robotic prey system

The robotic prey system enabled the prey to respond to an approaching predator by escaping once the predator had approached within a pre-specified distance of 27 cm. This system had five key components: a rectangular tank divided into an experimental arena and a holding zone (**Fig. S1a**), a Bluetooth-controlled robot on a wooden platform suspended below the experimental tank (**Fig. 1a**), an artificial prey item located within the experimental tank itself (**Fig. S1b**), a webcam positioned above the tank and a Bluetooth-enabled laptop connected to the webcam via a USB cable. During the test period of the experiment, blue acara cichlids left the refuge in 532 out of a total of 540 trials, and triggered the prey escape response in 524 of these trials (**Movies S1-S3**). There was no difference between the predictable and unpredictable treatments in the directional error of the escaping prey or in the prey’s reaction distance (**Table S1**).

### Predator behaviour during the approach phase

To investigate whether and how predators adjusted their approach behaviour in response to the prey escape strategy they encountered over successive trials, we compared a set of models predicting the maximum speed of the predator during the approach phase, i.e. the period before the prey escape response was triggered. Based on a comparison of AICc values, with a difference of two units or more indicating strong support for one model over another (40), the model including the interaction between treatment and the prey’s escape angle received most support from the data (**Table 1**). In the predictable treatment, the predators reached higher maximum speeds when approaching prey which escaped directly away from them (**Movie S1**) compared to prey that escaped at an acute angle (**Movie S2**), but in the unpredictable treatment, no relationship between maximum approach speed and prey escape angle was observed (**Fig. 2a**). The positive relationship between maximum approach speed and prey escape angle in the predictable treatment was not explained by differences between individual predators in traits which could influence approach speeds, such as body size or a proxy for the predator’s motivation (**Table S2**). There was no evidence for an effect of trial number on the predator’s maximum approach speed, or an interaction between treatment and trial number (**Table 1**), suggesting that treatment had no influence on how the predator’s maximum approach speed increased with further experience after the training period. The effect of trial number varied considerably between individual predators (**Fig. S3**), as demonstrated by the large reduction in model fit when individual-level random slopes for trial number were removed from the top-supported model (ΔcAIC: 65.2).

**Table 1:**
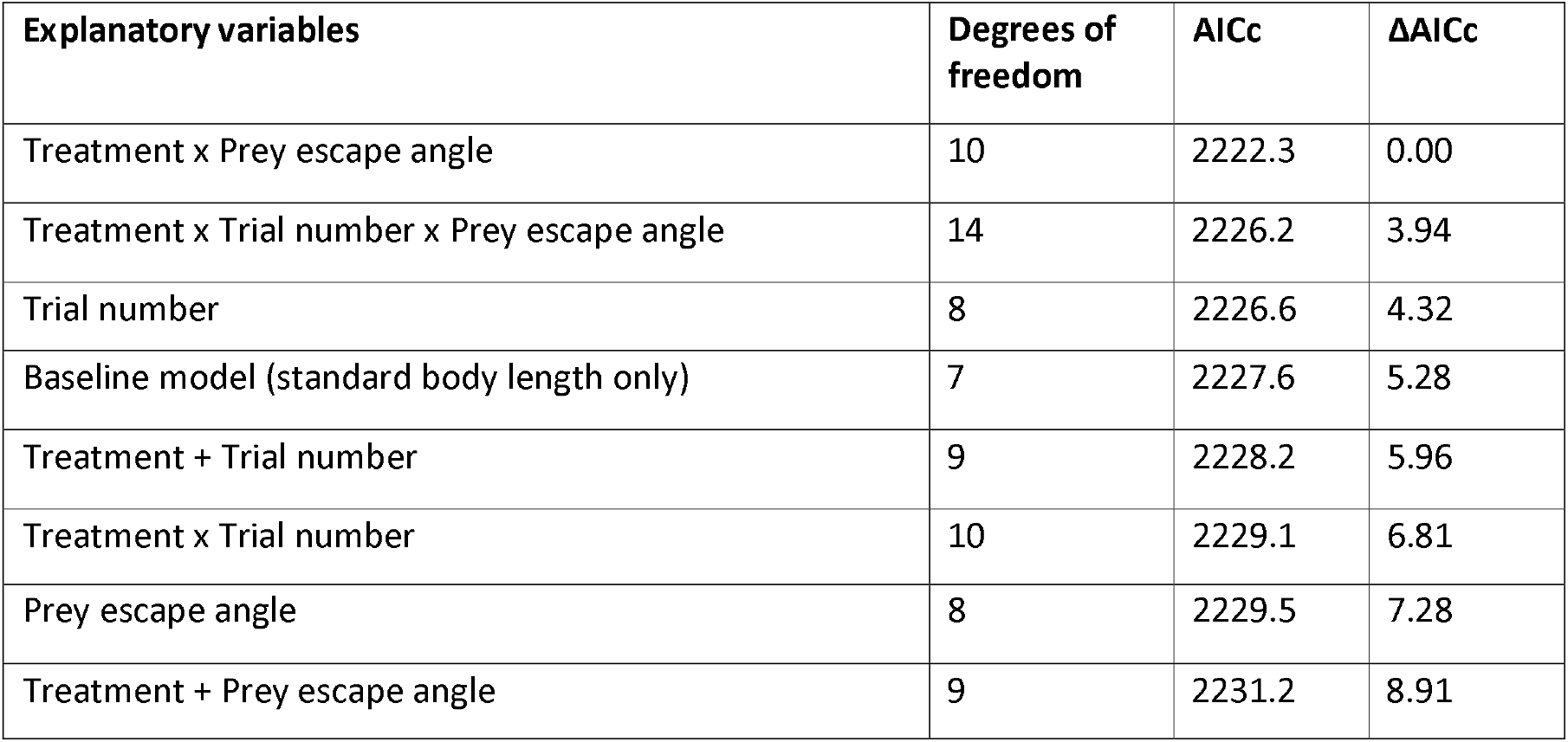
Results of LMMs (linear mixed-effects models) explaining variance in maximum speed of the predator during the approach phase. In order to ensure the predator was motivated to attack and pursue prey, this analysis was limited to the subset of 363 trials (featuring 25 individual predators) in which the predator approached the prey directly (i.e. at the time when the prey escape response was triggered, the bearing of the predator to the prey was less than 45°) and subsequently captured the prey. All models included standard body length as an explanatory variable, to control for differences in body size between individuals.

| Explanatory variables | Degrees of freedom | AICc | ΔAICc |
| --- | --- | --- | --- |
| Treatment x Prey escape angle | 10 | 2222.3 | 0.00 |
| Treatment x Trial number x Prey escape angle | 14 | 2226.2 | 3.94 |
| Trial number | 8 | 2226.6 | 4.32 |
| Baseline model (standard body length only) | 7 | 2227.6 | 5.28 |
| Treatment + Trial number | 9 | 2228.2 | 5.96 |
| Treatment x Trial number | 10 | 2229.1 | 6.81 |
| Prey escape angle | 8 | 2229.5 | 7.28 |
| Treatment + Prey escape angle | 9 | 2231.2 | 8.91 |

**Figure 2:**
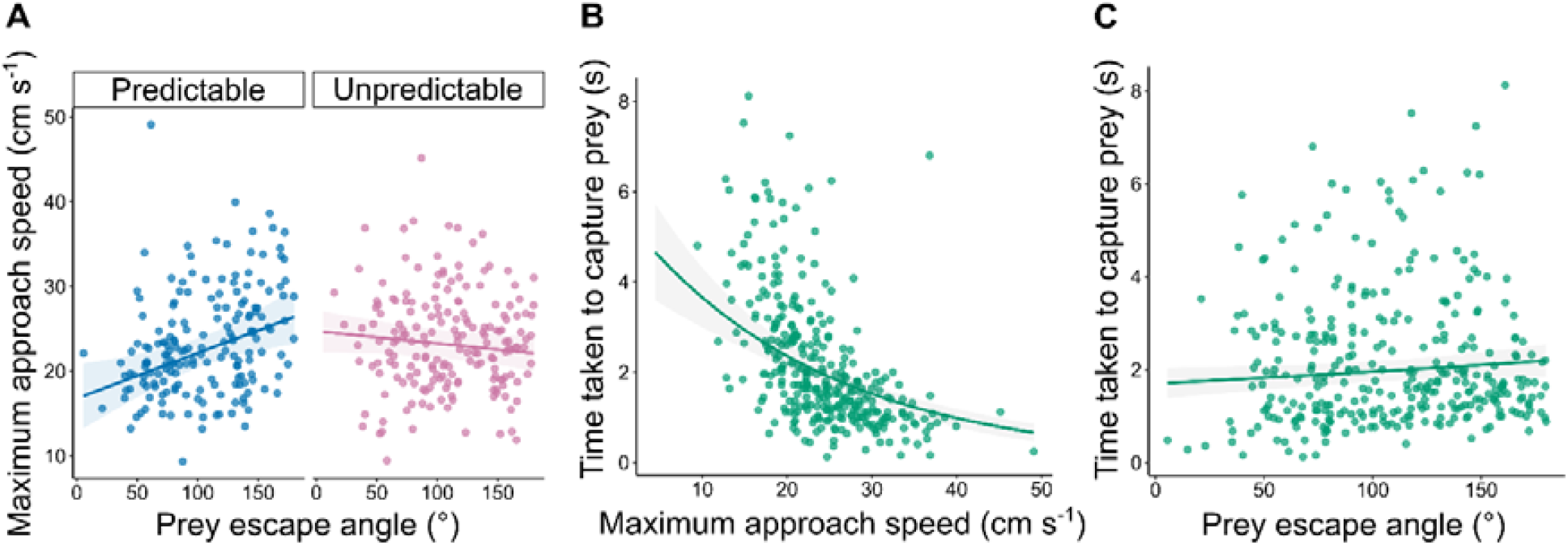
Determinants of the predator’s approach speed and the time taken to capture prey. (a) Influence of the interaction between treatment (predictable vs unpredictable) and prey escape angle on the maximum approach speed of the predator. (b) Effect of the predator’s maximum approach speed on the time taken to capture prey. (c) Effect of the prey’s escape angle on the time taken to capture prey. In (a-c), the coloured lines indicate the predicted fit from the top-supported model in **Table 1** (a) and **Table 2** (b, c), with other explanatory variables held constant at their mean values. The shaded area represents the 95% confidence intervals surrounding the predicted values. Each data point represents a single trial.

### Effects on the time taken to capture prey

In natural predator-prey encounters, where the prey can potentially flee to safety, the time taken by the predator to reach the prey is a key factor determining the outcome of the interaction (21). In our experiment, we therefore focused on the time taken to capture prey as the variable with greatest relevance to prey survival. The predator’s maximum approach speed had a large effect on the time taken to capture prey (the three models which included the predator’s maximum approach speed as an explanatory variable represented a substantial improvement in model fit over the baseline model featuring only reaction distance; **Table 2**), with faster approaches resulting in reduced latencies to capture (**Fig. 2b**). There was also evidence that increasing the prey’s escape angle increased the time taken to capture prey: the model including this main effect had the lowest AICc score by a margin of 0.65 units (**Table 2, Fig. 2c**). This can explain why predators in the predictable prey treatment reached faster maximum speeds when they approached prey expected to escape directly away from them, as a higher maximum speed could compensate for the additional time needed to capture prey fleeing directly away. This apparent compensation for longer capture times can also help clarify why the effect of treatment and prey escape angle on the predator’s maximum approach speed (the treatment × prey escape angle interaction shown in **Fig. 2a**) did not translate into an effect on the time taken to capture prey (**Table 2**).

**Table 2:** Results of Gamma GLMMs (generalised linear mixed-effects models) explaining variance in the time taken to capture prey. Since most approaches by the predator resulted in the prey being captured within less than 10 seconds (**Fig. S4**), trials in which fish failed to capture prey within this time limit were not considered in this analysis as the predator was unlikely to be sufficiently motivated to pursue prey in these trials, resulting in 325 observations of 23 individual fish. All models included reaction distance as an explanatory variable, to control for differences in the distance to the prey at the time when the prey escape response was triggered.

| Explanatory variables | Degrees of freedom | AICc | ΔAICc |
| --- | --- | --- | --- |
| Maximum predator approach speed + Prey escape angle | 9 | 791.8 | 0.00 |
| Maximum predator approach speed | 8 | 792.5 | 0.65 |
| Maximum predator approach speed x Prey escape angle | 10 | 793.9 | 2.08 |
| Prey escape angle | 8 | 839.7 | 47.83 |
| Baseline model (reaction distance only) | 7 | 839.7 | 47.87 |
| Treatment + Prey escape angle | 9 | 841.5 | 49.94 |
| Trial number | 8 | 841.8 | 49.97 |
| Treatment x Prey escape angle | 10 | 843.5 | 51.65 |
| Treatment + Trial number | 9 | 843.9 | 52.08 |
| Treatment x Trial number | 10 | 844.9 | 53.10 |
| Treatment x Trial number x Prey escape angle | 14 | 850.7 | 58.85 |

### Predator behaviour during the pursuit phase

The time to capture prey during the pursuit phase will depend on the speed, acceleration and manoeuvrability of the predator (39, 41). To examine the consequences of the prey’s predictability and escape angle, and the approach speed of the predator, on the kinematics of the pursuit, we considered data from the 117 trials in both treatments in which prey escaped at an acute angle (< 90°), where manoeuvrability would be most important. Predators which approached prey more rapidly reached higher maximum speeds during the subsequent pursuit, consistent with the faster capture times shown in **Fig. 2b**. After controlling for this effect, the maximum speed of the predator during the pursuit phase was faster if the prey escaped sideways (closer to 90°) than toward (closer to 45°) the predator (**Fig. 3a; Table 3**). There was no evidence for an effect of the prey’s predictability (treatment) on the maximum pursuit speed, or for an interaction between treatment and the predator’s maximum approach speed, which would be expected if the effects of a rapid approach depended on the prey’s predictability (**Table 3**). There was also no indication that the minimum speed of the predator during the first half of the pursuit was influenced by the predator’s maximum approach speed, the prey’s predictability or the interaction between these two variables (**Table S3**).

**Table 3:**
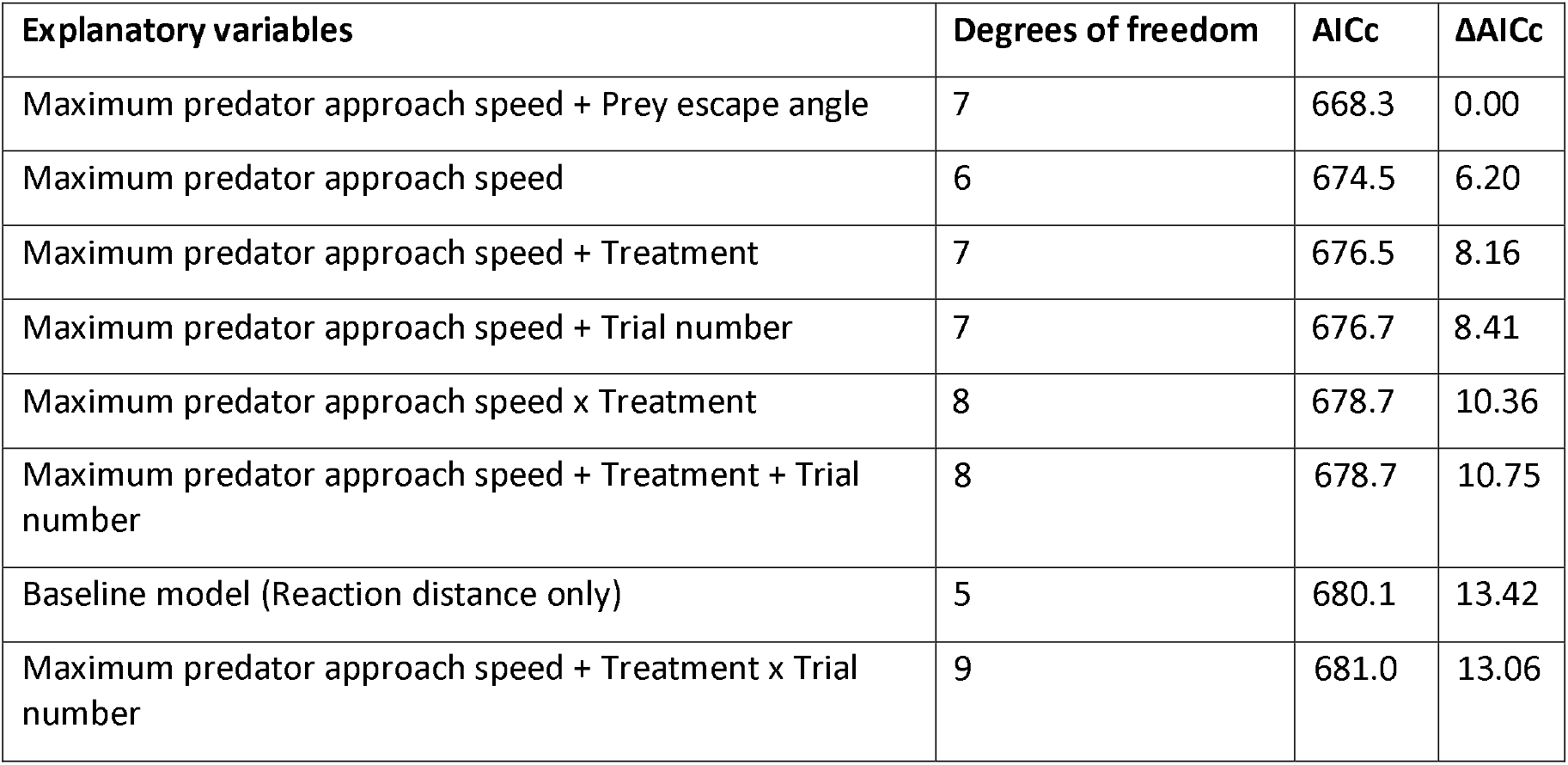
Results of LMMs explaining the variation in the maximum speed of the predator over the course of the pursuit, based on 117 observations of 19 individual fish in trials where prey escaped at an acute angle (< 90°). All models included reaction distance as an explanatory variable, to control for the effect of proximity to the prey at the point when the prey started to escape on changes in the predator’s speed. Maximum predator approach speed was also included in all models except the baseline (reaction distance only) model, to control for the expected effect of approach speeds on the maximum pursuit speed.

**Figure 3:**
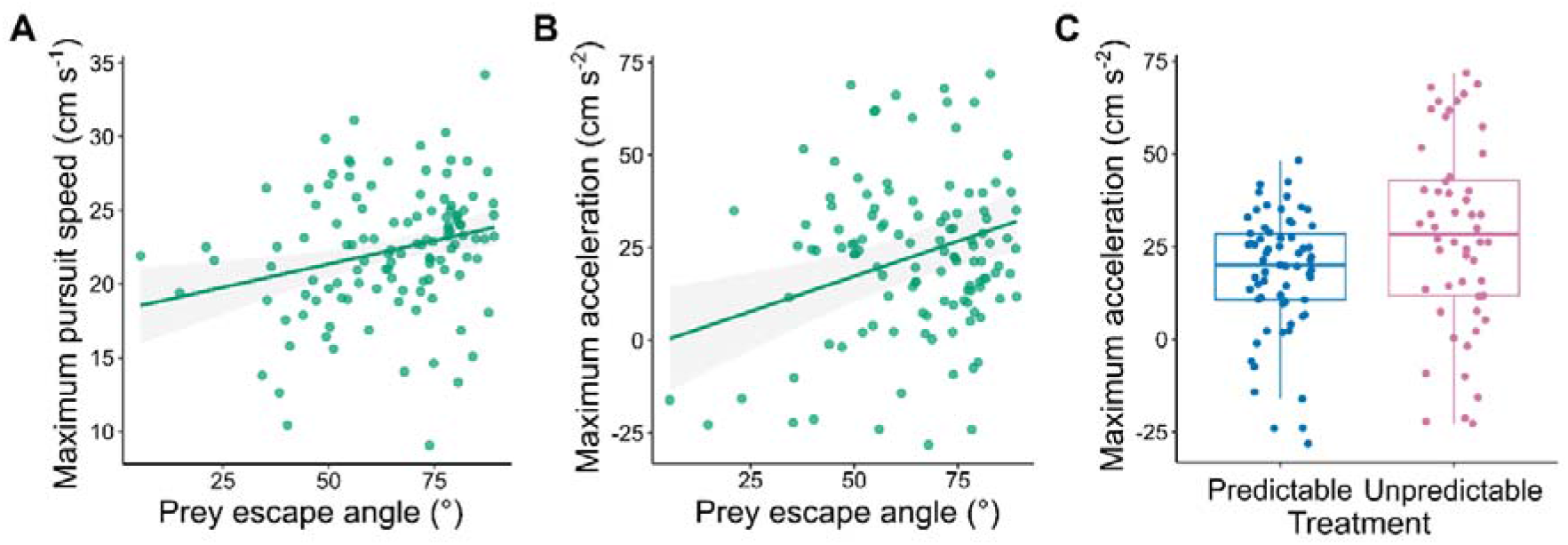
Behaviour of the predator during the pursuit of the robotic prey. Shown are the effects of prey escape angle (a, b) or treatment (c) on the maximum pursuit speed (a) and maximum acceleration (b, c) of the predator. In panels (a) and (b), the lines indicate the predicted fit from the top-supported model in **Tables 3** and **4**, respectively, with other explanatory variables held constant at their mean values. The shaded area represents the 95% confidence intervals surrounding the model fit. Each data point represents a single trial. Some prey escape angles were under 45° as the executed escape angles did not perfectly match the programmed escape angles.

The faster maximum speed of predators attacking prey with higher prey escape angles, i.e. those closer to 90°, can be explained by greater maximum accelerations when pursuing these prey (**Table 4; Fig. 3b**). There was also evidence for an effect of treatment on the maximum acceleration of the predator, as the model featuring maximum predator approach speed, reaction distance and treatment as explanatory variables received more support from the data than the model including only maximum predator approach speed and reaction distance (**Table 4**). The maximum acceleration when pursuing prey escaping in an unpredictable direction was higher than during the pursuit of predictable prey (**Fig. 3c**). Increased approach speeds were also associated with greater maximum deceleration during the first half of the pursuit (**Table S4**), as would be expected following a quick approach. However, the predator’s turning during the pursuit phase was unrelated to the predator’s maximum approach speed or treatment (**Table S5** and **S6**).

**Table 4:** Results of LMMs explaining variation in the maximum acceleration of the predator throughout the pursuit, based on 117 observations of 19 individual fish in trials where prey escaped at an acute angle (< 90°). All models included reaction distance as a explanatory variable, to control for the effect of proximity to the prey at the point when the prey started to escape on changes in the predator’s speed. Maximum predator approach speed was also included in all models except the baseline (reaction distance only) model, to control for the expected effect of approach speeds on maximum acceleration in the pursuit.

| Explanatory variables | Degrees of freedom | AICc | $\Delta AICc$ |
| --- | --- | --- | --- |
| Maximum predator approach speed + Prey escape angle | 7 | 1030.9 | 0.00 |
| Maximum predator approach speed + Treatment | 7 | 1037.7 | 6.77 |
| Baseline model (Reaction distance only) | 5 | 1038.0 | 7.07 |
| Maximum predator approach speed + Treatment x Trial number | 9 | 1038.7 | 7.77 |
| Maximum predator approach speed x Treatment | 8 | 1038.8 | 7.89 |
| Maximum predator approach speed | 6 | 1039.6 | 8.72 |
| Maximum predator approach speed + Treatment + Trial number | 8 | 1040.3 | 9.37 |
| Maximum predator approach speed + Trial number | 7 | 1041.9 | 10.98 |

## Discussion

The potential antipredator benefits of behaving unpredictably have long been recognised (6, 42, 43), yet it remained unclear how real predators respond to unpredictable escape tactics in their prey. By developing a novel experimental system in which robot-controlled prey fled from a blue acara cichlid predator, we manipulated whether each individual predator experienced predictable prey that always escaped at the same angle relative to the predator’s approach, or whether this angle varied unpredictably from trial to trial. Our results demonstrate that predators adjust their approach behaviour (their speed) when they are able to predict the direction in which prey will escape. Prey escaping directly away from the predators (closer to 180°) were approached more quickly even before the prey responded, which compensated for the longer time needed to capture such prey during the pursuit, compared to those escaping at an acute angle. If the predators attempted to minimise the time to capture prey, however, they should have approached prey expected to escape at acute angles just as rapidly, given that the benefit of a rapid approach applied equally across all prey escape angles. When examining the kinematics of the predator’s trajectory during the pursuit, even after controlling for the faster approach speed, the predators accelerated more and reached faster maximum speeds when pursuing prey fleeing at angles closer to 90° than 45°. These results suggest that with information on the prey’s likely escape direction, the predators were minimising the costs of capturing prey and aimed to achieve an adequate, rather than minimal, time to capture.

Confirming that predators in the unpredictable treatment could not anticipate the prey’s escape direction, there was no relationship between the predator’s approach speed and the escape angle of unpredictable prey. Instead, these predators adopted intermediate approach speeds, which may represent a form of insurance aimed at buffering against the uncertainty caused by a lack of consistency in the prey’s escape direction (44). As approach speeds were a key determinant of the time taken to capture prey during the pursuit, intermediate approach speeds in the unpredictable treatment trials resulted in no effect of the prey’s predictability on average capture times. These results provide further evidence that the predators were maintaining an adequate level of performance rather than minimising the time taken to capture prey.

In our study, the variable most directly relevant to prey survival was the time taken to capture prey, as a longer delay increases the probability that the prey can find refuge or the predator gives up the pursuit (45-47). We did not observe an effect of prey predictability on capture times, either as a main effect or as part of an interaction, bringing into question whether unpredictable escape behaviour has an adaptive benefit to prey by increasing the chances of survival. Instead, the unpredictable behaviour observed in real prey (20-25) may come about through biomechanical or sensory constraints on the angle of escape (26). However, when faced with prey escaping at an acute angle, the predators accelerated more when pursuing unpredictable compared to predictable prey, which could result in the accumulation of substantial energetic costs for predators over longer pursuits where prey repeatly change their direction unpredictably (41, 48, 49). If there is a greater cost of preying upon unpredictable prey, this may result in predators switching to other prey, either to an alternative target nearby (for prey in groups) or by searching for and choosing to pursue other prey types (50, 51).

In their natural environment, blue acaras are opportunistic predators which actively pursue prey such as guppies (36, 52). The results of our study are relevant to predator-prey interactions involving pursuit predators like blue acaras which adjust their trajectory in response to fleeing prey (53, 54), rather than those involving ambush or stalk-and-strike predators that attack prey at close range and do not adjust their trajectories in the period immediately after striking (55, 56). In the latter case where predators have a limited capacity to adjust, unpredictable prey escape behaviour may be more effective than in our study. Our results underscore the importance of considering predators as responsive participants within predator-prey interactions (57, 58), capable of flexibly adjusting their behaviour depending on the prey’s tactics.

## Materials and Methods

### Experimental subjects and housing

A total of twenty-eight blue acara cichlids (*Andinoacara pulcher*) were tested in the experiment (median standard body length: 6.2 cm, inter-quartile range: 1.95 cm). As a maximum of 16 fish could be tested simultaneously, the experiment was first conducted with 16 fish in November and December 2018 and then repeated with an additional 12 fish in February and March 2019. Outside of the experimental period, fish were kept in glass tanks (width = 40 cm, length = 70.5 cm, height = 35.5 cm), with a daily 12h:12h dark:light cycle. The water temperature inside the tanks was maintained at 26-27°C (+/- 0.5°C). Throughout the training and test periods of the experiment, groups of four fish were kept in a holding zone located at one end of the experimental arena. Groups comprised individual fish of different sizes, enabling individuals to be identified. Throughout the experiment, fish were fed *ad libitum* on aquarium fish pellets (ZM Systems, Large Premium Granular) at the end of each day.

### Experimental set-up and robotic prey system

Trials were filmed using a camcorder (Panasonic SD 800, resolution: 1920 x 1080 pixels, frame rate: 25 frames per second) suspended 225 cm above the experimental tank. A webcam (Logitech C920 USB Pro) was also secured 175 cm above the tank to monitor the movements of the predator in real time during experimental trials. The Bluetooth-controlled robot (MiaBot PRO BT v2, Merlin Systems Corp. Ltd.) consisted of two wheels set within either side of a 7.5 cm cube containing the electronic circuitry, batteries and separate motors for each wheel. The artificial prey item was a small amount of food (an approximately 5 mm x 8 mm piece of defrosted fish) attached to a length of transparent monofilament fibre (thickness = 1 mm, length = 4 cm) protruding from a cone-shaped white plastic base (diameter = 2.5 cm, cone height = 1.8 cm, **Fig. S1b**). Movement of the prey was controlled by the robot through a connection between magnets embedded in the base of the prey item, and another magnet embedded in a plastic hood secured to the top of the robot. In training or test trials in which the prey was programmed to respond to an approaching predator, the artificial prey item was placed in the same starting position within the experimental arena (**Fig. S1a**). Detection of the approaching predator was integrated with control of the robot’s movements using a custom-built program (**Fig. S2**), written in python (version: 2.7.12), and utilising the OpenCV library (version: 3.1.0). By processing footage captured by the webcam positioned above the experimental arena, the program triggered the prey’s escape response by sending movement commands to the robot once the predator had approached within 27 cm of the prey’s starting position. In all escape responses, the robot was programmed to move a total of 38.5 cm in a straight line from its starting position towards the periphery of the tank. Only the prey’s initial escape angle, measured relative to the approaching predator, was manipulated, with all other parameters held constant. The turning time of the robot was standardised across trials by including a 0.3 second time delay within the program, enforcing a consistent time lag between the initial turn command being sent to the robot via Bluetooth and the subsequent movement command which directed the robot to escape in a straight line. Pilot trials filmed at a frame rate of 240 frames per second (GoPro Hero 5, resolution: 1280 x 720 pixels) indicated that the mean overall time delay between the predator moving within range and the robot moving was 0.7 seconds (n = 7, standard deviation: 0.08 seconds).

### Training period

Before being trained to attack artificial prey, the boldness of each individual fish was measured by recording the time taken to leave the refuge and enter the experimental arena, in two separate trials conducted 2 days apart (data from these trials was not used further in this study). As individuals in groups tend to behave more boldly than lone individuals (59), groups of fish were subsequently progressively trained to approach and take food from the artificial prey item in a series of training trials (further details are provided in the **Supplementary Text**).

### Test period

After completing the training period, each fish was tested individually in twenty successive experimental trials with either predictable or unpredictable prey (**Fig. 1c**). At the start of each 10-minute experimental trial, fish were transferred to the central refuge and left to habituate for 3 minutes. After 3 minutes, the sliding door was opened, allowing fish to enter the experimental arena.

Experimental trials took place in three six-day blocks and one final two-day block, with a one-day gap occurring after each block. Trials took place between 0900 and 1700, and the order individuals were tested in was randomised on each day. Individual fish were allocated randomly to either the predictable or the unpredictable treatment, subject to the constraint that every group of four fish from the same holding compartment was split equally between the two treatments, with the largest two fish in each group being assigned to different treatments. In the unpredictable treatment, escape angles were generated randomly in each trial, and in the predictable treatment, trials with the same individual predator were conducted with a single randomly generated escape angle. As in the training period, in both treatments, escape angles were chosen from a uniform distribution from 45° to 315° (where 0° was defined as the approach angle of the predator). Throughout all trials in the test period, the robot was programmed to escape at a speed of 15.8 cm s ^-1^.

### Video analysis

ToxTrac (version 2.84) was used to extract the position of the predator in each video frame up to 30 seconds before and 30 seconds after the prey escape response was triggered (60). The coordinates of the escaping prey were extracted manually from each frame using a custom-built program written in python (version: 3.6.9) using the OpenCV library (version: 4.1.1). Multiple behavioural variables were also manually extracted from videos, including the time taken for the predator to leave the refuge, relative to the door opening. The time taken to trigger the prey escape response, relative to the predator’s emergence from the refuge, was also calculated as a measure of the predator’s motivation to pursue prey within each trial. Additionally, the time taken to capture prey was defined as the time difference between the moment the prey started to escape and the point at which the predator made physical contact with the prey.

R version 3.5.1 was used to calculate all predator movement variables from the raw positional data (61). Predator and prey trajectories were combined to calculate the predator’s bearing to the prey, which was defined as the absolute angular difference between the predator’s heading and the straight line connecting the positions of the predator and the prey. Since spurious changes in heading might result from tracking error when the predator is stationary, heading angles were only calculated when the predator had moved a minimum distance of 0.5 cm between frames. Throughout the analysis, the prey escape angle was defined relative to the approaching predator, ranging from approximately 45° to 180°, with 180° indicating that the prey had escaped directly away from the predator (**Fig. 1b**). As the actual prey escape angle sometimes deviated from the programmed escape path, all statistical analyses were based on realised escape angles, calculated from the known start and end points of the prey’s escape trajectory. Additionally, although the prey was programmed to respond when the predator had approached within 27 cm, the actual prey reaction distance varied due to a combination of a short delay between the predator being detected and the initial movement of the prey and differences in predator approach speeds across trials. Reaction distances were therefore calculated as the distance between the predator and prey positions in the video frame immediately before the escape response was initiated.

In the period before the prey escape response was triggered, predators tended to accelerate in a straight line towards prey during their approach. As the only information available to the predator during the approach phase about the prey’s subsequent escape was from its experience of previous trials, the predator’s maximum approach speed in each trial was calculated to measure how prey predictability influenced predator behaviour. To obtain reliable estimates of maximum approach speeds and reduce noise, LOESS (locally weighted regression) with a smoother span of 0.1 was used to smooth raw speed values over time. To avoid generating a smoothed time series which includes negative values, which were sometimes produced when using LOESS to smooth data on the original scale, the raw speed values were log(x+1) transformed prior to smoothing. Smoothed speeds on the original scale were then obtained by applying the inverse of this transformation, resulting in positive smoothed speed values. The maximum predator approach speed was defined as the highest speed during the period of the approach phase in which the predator headed continuously towards the prey (i.e. the predator’s bearing to the prey did not exceed 45°), and did not subsequently deviate from this overall direction by heading away from the target. The period up to 10 seconds before the prey started to escape was considered. Data from trials when predators triggered the prey escape response while approaching at a bearing of greater than 45° to the prey were not included in the analysis, as fish were unlikely to have been as motivated to attack in these trials.

During the pursuit phase (the period after the prey escape response was triggered), the maximum and minimum speeds and acceleration of the predator were derived from smoothed speed values, obtained using the same LOESS method described above. To quantify the predator’s turning performance, both the predator’s maximum turn speed and minimum turn radius were also calculated, providing an indication of how rapidly and how sharply the fish turned during the pursuit (62). Turn speed was defined as the change in the direction of the predator’s heading in successive frames, and turn radius was calculated as the straight-line distance between the predator’s position in frames *i* and *i*+ 2, divided by two times the sine of the change in the predator’s heading *θ* between successive frames, *i* and *i*+ 1 (below, x and y indicate the x- and y-coordinates of the fish):

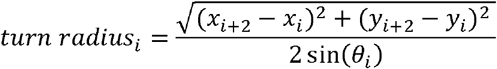

### Statistical analysis

R version 3.5.1 was used to conduct all statistical analyses (61). Both linear mixed effects models (LMMs, fitted with the lme4 package) and generalised linear mixed effects models (GLMMs, fitted with the glmmTMB package) were used to explore the variables that impacted the behaviour of the predators in the approach and pursuit phases. Throughout the analysis, the relative influence of explanatory variables was assessed using AICc (Akaike’s Information Criterion corrected for small sample sizes) values to compare the level of support from the data for a particular model within a set of candidate models. To limit the number of candidate models being compared, simpler versions of three-way interaction models (models lacking the three-way interaction but retaining the constituent two-way interactions) were not included in the initial model comparison set (40, 63). These were only considered if the initial model comparison revealed that the three-way interaction had an important effect. Within model comparison sets, additional explanatory variables were also added to all models in order to account for potentially confounding effects such as the standard body length of the predator, reaction distance (used to control for differences in the distance to the prey when the prey escape response was triggered) and the predator’s maximum approach speed (used to control for the expected effect of approach speed on pursuit speed or acceleration). An additional model featuring only these control variables was also included in each model comparison set to serve as the baseline for comparison with the other models. Where control variables were also of interest, a null model lacking any explanatory variables was also included.

Within each comparison set, all models shared the same random effects structure. To control for similar experiences during the training period, and to account for repeated measures of the same individuals, random intercepts for training group and individual identity were included in all models. Individual-level random slopes for the effect of trial number were also included in models with maximum predator approach speed and the time taken to capture prey as the response (64). The extent of inter-individual differences in the effect of trial number on maximum approach speed was also assessed by comparing the conditional AIC (cAIC) of an LMM with individual-level random slopes to an otherwise identical model lacking random slopes using the cAIC4 package in R (version: 0.9; 65). To aid model fitting, continuous explanatory variables were scaled prior to being included in the model, by subtracting the mean and dividing by the standard deviation. Model assumptions were checked by examining QQ-plots of the residuals, residuals vs. fitted values and the distribution of the conditional modes of the random intercepts. The DHARMa package (version: 0.2.7) was used to check model assumptions for GLMMs (66).

## Acknowledgements

The authors would like to thank Peter Gardiner at the University of Bristol Animal Services Unit for his help caring for the fish, and Alexander Campo at Université Libre de Bruxelles for advice on how to program the robots.

