## Supplementary figures and tables for "Responsive robotic prey reveal how predators adapt to predictability in escape tactics"

**Contents of Supplementary Material**

Supplementary Text: Description of training period methods.

Fig. S1: Components of the robot prey experimental system.

Fig. S2: Schematic diagram showing steps involved in the program used to control how prey respond to an approaching predator.

Fig. S3: The relationship between maximum approach speed and trial number, for each individual fish tested in the experiment.

Fig. S4: The distribution of the time taken by the predator to capture prey.

Table S1: Analysing the performance of the robotic prey system.

Table S2: Results of LMMs explaining variance in maximum speed of the predator during the approach phase, in the predictable treatment, based on 179 observations of 12 individual fish.

Table S3: Results of LMMs explaining the variation in the minimum speed of the predator over the course of the pursuit, based on 117 observations of 19 individual fish in trials where prey escaped at an acute angle (< 90°).

Table S4: Results of LMMs explaining variation in the minimum acceleration of the predator, over the first half of the pursuit, based on 117 observations of 19 individual fish in trials where prey escaped at an acute angle (< 90°).

Table S5: Results of Gamma GLMMs explaining variation in the maximum turning speed of the predator during the pursuit phase, based on 116 observations of 19 individual fish in trials where prey escaped at an acute angle (< 90°).

Table S6: Results of LMMs explaining variation in the minimum turn radius of the predator during the pursuit phase, based on 115 observations of 19 individual fish in trials where prey escaped at an acute angle (< 90°).

Movie S1: Representative example of a predictable treatment trial, with the prey escaping directly away from the predator (close to 180°).

Movie S2: Representative example of a predictable treatment trial, with the prey escaping directly at an acute angle from the predator (< 90°).

Movie S2: Representative example of an unpredictable treatment trial, showing the predator accelerating towards the prey during the pursuit phase as the prey escapes.

**Supplementary Text:** Description of training period methods

During the training period, groups of fish were progressively trained to approach and take food from the artificial prey item in a series of training trials with three sequential training stages. Training trials were conducted in the same groups that the fish were housed in. Groups progressed to the next training stage once a pre-specified criterion was reached, ensuring all groups were trained to a similar level. Before each training trial, groups of fish were transferred to the central refuge and left to habituate for 3 minutes. After 3 minutes, the sliding door was opened, allowing fish to enter the experimental arena. Training trials lasted 10 minutes. At each stage of the training process, groups were subjected to three training trials per day.

In the first stage of the training period, a static, baited prey item was positioned 25 cm from the entrance of the refuge, surrounded by a small amount of food (12 cichlid pellets). Successful trials were those in which at least one fish from the group consumed a pellet and progression to the next stage occurred after success in 6 consecutive training trials. Training trials in the second stage were identical to those in the first stage, but the prey item was placed at the centre of the experimental arena, in the same location as the test trials. In stage two (and stage three), successful trials were those where at least one fish from the group consumed the food attached to the artificial prey item. Success in 10 consecutive trials was required to progress to the third (final) stage of the training period. The baited, robot-controlled prey item was programmed to initiate an escape response when a fish approached within the set distance (27 cm) at a speed set at 7.9 cm s^-1^ (half the speed used in experimental trials). To ensure that the training process did not bias the response of fish towards either of the two experimental treatments, during the third training stage groups were exposed to prey which escaped consistently in the same direction, as well as prey which escaped at a random angle. Within each pair of consecutive training trials (i.e. the pair formed by the first and second training trials, third and fourth trials, fifth and sixth trials, etc.), groups of fish experienced one trial with consistently escaping prey and one trial with randomly escaping prey. Within a pair of training trials, the order of consistent and random trials within each pair of trials was randomised for each group of fish. This way of determining trial order was repeated within pairs of consecutive training trials, until the fish had reached the criterion for progression to the test period. The prey’s escape angle during consistent trials was chosen separately for each group of fish. To prevent prey moving directly back towards an attacking predator, in both consistent and random trials, escape angles were chosen from a uniform distribution ranging from 45° to 315° (where 0° was defined as the approach angle of the predator; **Fig. 1b**).

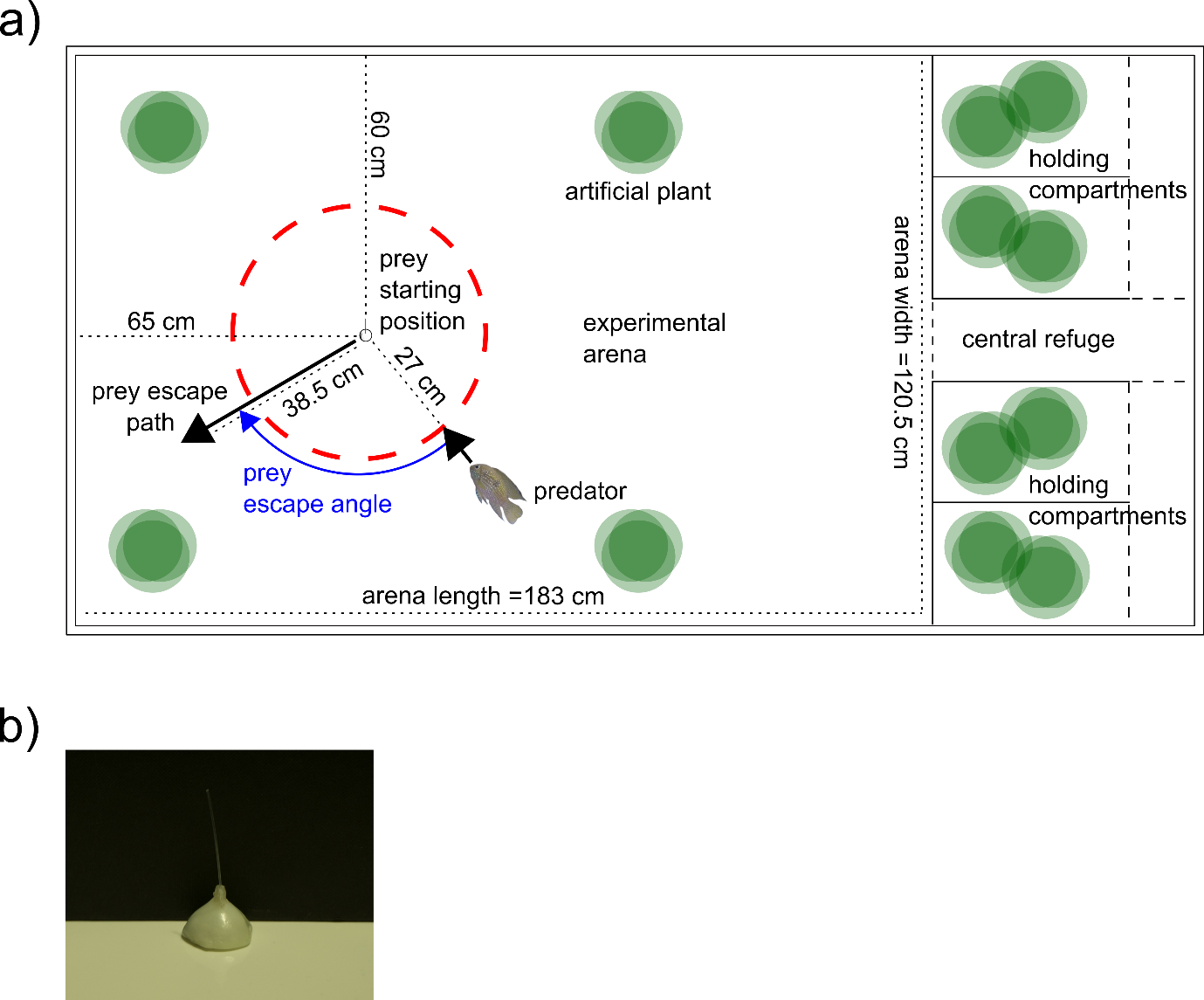

**Fig. S1:** Components of the robot prey experimental system. (a) Scale diagram of the experimental arena, viewed from above. The red dashed line indicates the predator-prey distance (27 cm) at which the initially stationary prey item was programmed to escape from an approaching predator. Black arrows indicate the heading of the approaching predator and the escaping prey. The prey escape angle (relative to the approaching predator) is shown in blue. The experimental arena and fish holding zone were situated within a large rectangular aluminium tank (width = 127 cm, height = 36 cm, length = 238 cm). Within this external structure, an inner tank was constructed from white PVC walls (height = 35 cm, thickness = 0.8 cm) sealed to a white base made of compressed white foamed PVC (thickness = 0.2 cm) using aquarium sealant. This created a high contrast background enabling the movements of the predator to be tracked. The inner tank was divided into a large rectangular experimental arena, separated from a smaller holding zone by a white plastic divider, positioned at one end of the tank. The holding zone was further sub-divided into four compartments (width = 25.75 cm, length = 41.5 cm each), with two compartments positioned either side of a central refuge (width = 16 cm, length = 55.5 cm), which was covered by rigid plastic mesh. Each compartment contained a cylindrical tube and two artificial plants to provide cover for the fish. Small pebbles (< 0.5mm diameter) were also scattered across the floor of each compartment and the central refuge. Holding compartments were linked to the central refuge via a connecting corridor (width = 14 cm, length = 51.5 cm), which bordered the external wall of the inner tank. The holding compartments were separated from the central refuge and experimental arena by retractable doors (indicated by black dashed lines), enabling fish to be transferred from their holding compartments to the central refuge or released into the experimental arena without being caught in nets, thus minimising handling stress. The experimental arena also contained a water heater attached to the wall furthest from the holding zone, and four artificial plants positioned near the edges. Throughout the experiment, the water depth in the experimental tank was kept at 15 cm, temperature was held constant at 27°C (+/- 0.5°C) and a 12h:12h light:dark cycle was maintained. Water was continuously filtered and circulated throughout the entire tank using two Eheim Classic 600 external canister filters, which took in water via inflow pipes positioned in the corners of the arena closest to the heater and discharged water back into the holding zone. (b) Photo of the artificial prey item used in the experiment, shown against a dark background.

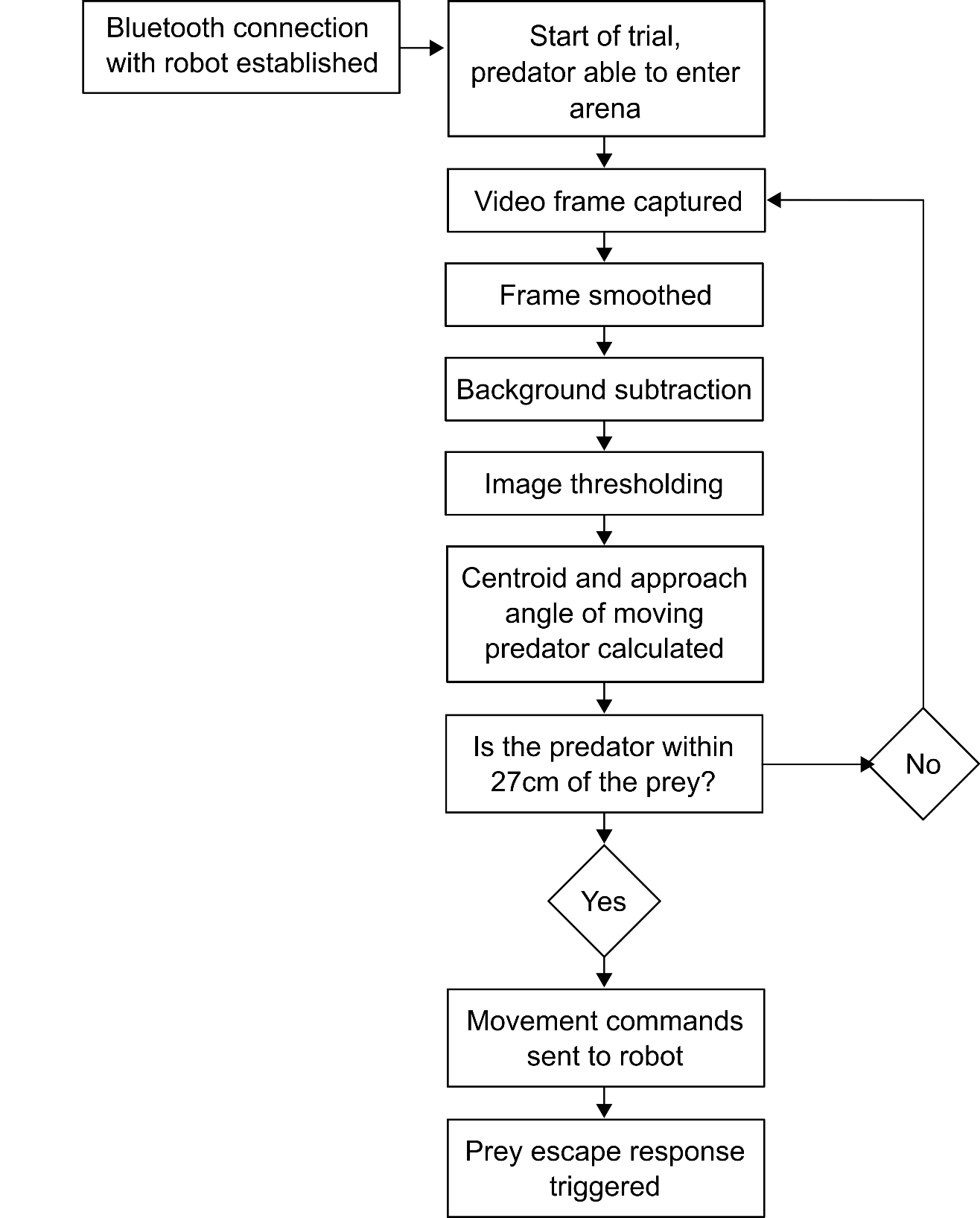

**Fig. S2:** Schematic diagram showing steps involved in the program used to control how prey respond to an approaching predator. Prior to an attack, the program continuously analysed video frames captured by the webcam positioned above the arena, monitoring any changes from one frame to the next which could indicate movement of the fish within the arena. As part of this process, each frame was first converted to grayscale and smoothed to filter out noise. Background subtraction was then used to calculate differences in pixel values between the current frame and a representation of the static background (i.e. the unchanging aspects of the experimental arena). The background was estimated using a running average, in which motion during more recent frames was weighted more heavily. Thresholding was then applied to the resulting image to isolate areas of the current frame which differed substantially from the background. Additional size filtering also ensured that any regions of movement below a pre-specified size threshold were disregarded, to make certain that noise was ignored and that the sole region of detected motion corresponded to the predator. The program then used this information to calculate the centroid of the predator in each frame, until the predator approached within a pre-specified radius of the prey’s starting position (27 cm in both training and test period trials). Video frame processing was halted at this point, and movement commands were sent to the robot, based on the predator’s angle of approach and the programmed prey escape angle, in order to execute the prey’s escape response.

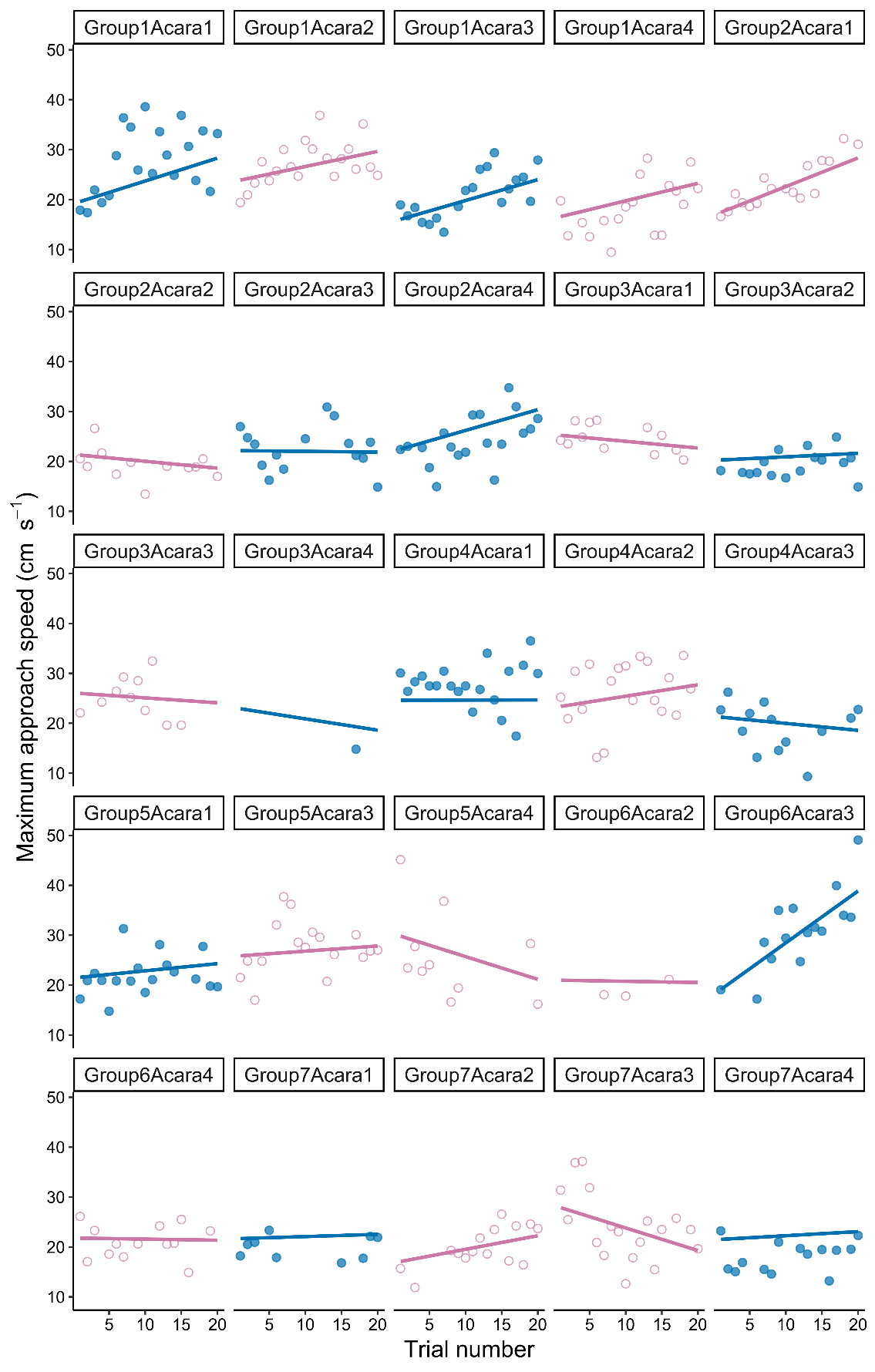

**Fig. S3:** The relationship between maximum approach speed and trial number, for each individual fish tested in the experiment. Colours indicate which treatment an individual was assigned to (predictable: solid blue points; unpredictable: empty pink points). Individual predicted slopes were estimated using the *predict* function in the R package lme4.

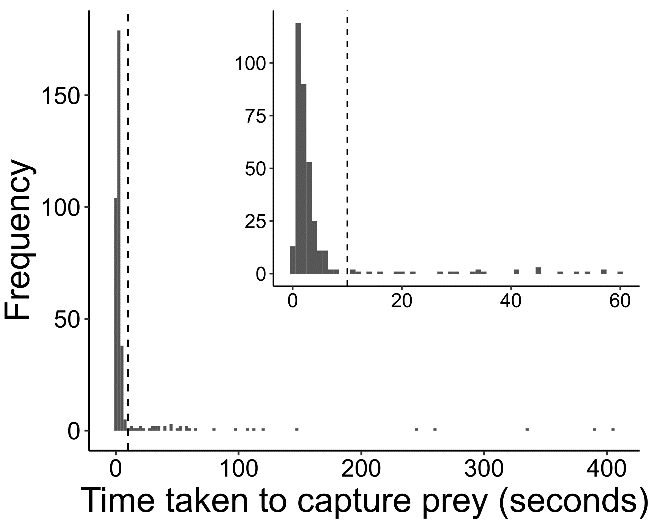

**Fig. S4:** The distribution of the time taken by the predator to capture prey. Data is shown for all 363 trials in which the fish left the refuge, and captured prey following a direct approach (i.e. trials in which the bearing of the predator to the prey was less than 45°, at the time when the prey escape response was triggered). The inset shows a magnified version of the same histogram, ranging from 0 to 60 seconds. In both the main histogram and the inset, the dashed vertical line indicates the 10 second threshold used to remove trials in which fish were insufficiently motivated to pursue the prey.

**Table S1:** Analysing the performance of the robotic prey system. The table compares AICc scores for Gamma GLMs (generalised linear models) constructed to explain variation in the angular difference between expected and realised prey escape angles, and for LMMs (linear mixed-effects models) explaining variation in reaction distance. Throughout the experiment, the programmed (i.e. expected) prey escape angle did not always perfectly match the escape angle that was realised: the median angular difference between the expected and realised prey escape angle was 8.5° (inter-quartile range, IQR = 10.7°). The directional error of the robotic prey system did not differ between the predictable and unpredictable treatments and was not correlated with the expected prey escape angle (as demonstrated by the lack of improvement in fit for models featuring these explanatory variables, compared to the null model). Additionally, although the realised predator-prey distance at which the prey initiated its escape response (reaction distance) varied from trial to trial, there was no overall difference in reaction distance between the two treatments.

| **Response variable** | **Explanatory variables** | **Degrees of freedom** | **AICc** | **ΔAICc** |
| --- | --- | --- | --- | --- |
| Angular difference between expected and realised escape angles | Treatment | 3 | 3773.6 | 0.00 |
|  | Null model (no explanatory variables) | 2 | 3775.6 | 1.97 |
|  | Expected escape angle | 3 | 3776.2 | 2.62 |
| Reaction distance | Null model (no explanatory variables) | 3 | 2281.3 | 0.00 |
|  | Treatment | 4 | 2282.0 | 0.73 |

**Table S2:** Results of LMMs explaining variance in maximum speed of the predator during the approach phase, in the predictable treatment, based on 179 observations of 12 individual fish. In the predictable treatment, individual predators were always exposed to prey escaping at the same angle. To test whether the positive relationship between the prey’s escape angle and the predator’s maximum approach speed (in predictable treatment trials; **Fig. 1c**) did not arise because of differences between individual fish, we examined which variables predicted the predator’s approach speed. The model including prey escape angle received most support from the data, compared to models featuring standard body length, the approach speed of the predator in the first trial or the time taken to trigger the prey escape response (an indicator of the predator’s motivation within a trial). This suggests that the relationship between maximum approach speed and prey escape angle observed in the predictable treatment (**Fig. 1c**) was unlikely to have arisen due to inter-individual variation in speed, body size or motivation.

| **Explanatory variables** | **Degrees of freedom** | **AICc** | **ΔAICc** |
| --- | --- | --- | --- |
| Prey escape angle | 7 | 1082.0 | 0.00 |
| Maximum approach speed in first trial | 7 | 1087.0 | 4.97 |
| Time taken to trigger the prey escape response | 7 | 1087.7 | 5.67 |
| Trial number | 7 | 1088.0 | 6.03 |
| Standard body length | 7 | 1089.0 | 6.95 |
| Null model (no explanatory variables) | 6 | 1089.5 | 7.51 |

**Table S3:** Results of LMMs explaining the variation in the minimum speed of the predator, during the first half of the pursuit phase, based on 117 observations of 19 individual fish in trials where prey escaped at an acute angle (< 90°). This first half of the pursuit was defined as period from when the prey started moving, until the time half-way between this start point and the moment the predator captured the prey. All models included reaction distance as a explanatory variable, to control for the effect of proximity to the prey at the point when the prey started to escape on changes in the predator’s speed. Maximum predator approach speed was also included in all models except the baseline (reaction distance only) model, to control for the expected effect of approach speeds on the minimum pursuit speed.

| **Explanatory variables** | **Degrees of freedom** | **AICc** | **ΔAICc** |
| --- | --- | --- | --- |
| Baseline model (Reaction distance only) | 5 | 725.5 | 0.00 |
| Maximum predator approach speed + Prey escape angle | 7 | 726.7 | 1.18 |
| Maximum predator approach speed | 6 | 727.1 | 1.57 |
| Maximum predator approach speed + Trial number | 7 | 727.3 | 1.80 |
| Maximum predator approach speed + Treatment | 7 | 728.2 | 2.68 |
| Maximum predator approach speed + Treatment + Trial number | 8 | 728.8 | 3.24 |
| Maximum predator approach speed x Treatment | 8 | 730.5 | 4.98 |
| Maximum predator approach speed + Treatment x Trial number | 9 | 731.0 | 5.44 |

**Table S4:** Results of LMMs explaining variation in the maximum deceleration of the predator, over the first half of the pursuit, based on 117 observations of 19 individual fish in trials where prey escaped at an acute angle (< 90°). This first half of the pursuit was defined as period from when the prey started moving, until the time half-way between this start point and the moment the predator captured the prey. All models included reaction distance as an explanatory variable, to control for the effect of proximity to the prey at the point when the prey started to escape on changes in the predator’s speed. Maximum predator approach speed was also included in all models except the baseline (reaction distance only) model, to control for the expected effect of approach speeds on the maximum deceleration in the pursuit. Compared to a baseline model including maximum predator approach speed and reaction distance as explanatory variables, only the model featuring trial number, maximum predator approach speed and reaction distance represented a substantial improvement. As the trials progressed, the fish decelerated more. The relatively large difference in AICc scores between the model including maximum predator approach speed and the baseline model indicated that the predator’s maximum approach speed was strongly negatively correlated with its maximum deceleration during the pursuit phase, consistent with greater deceleration by rapidly approaching fish.

| **Explanatory variables** | **Degrees of freedom** | **AICc** | **ΔAICc** |
| --- | --- | --- | --- |
| Maximum predator approach speed + Trial number | 7 | 969.8 | 0.00 |
| Maximum predator approach speed + Prey escape angle | 7 | 971.2 | 1.42 |
| Maximum predator approach speed + Treatment x Trial number | 9 | 971.9 | 2.07 |
| Maximum predator approach speed + Treatment + Trial number | 8 | 972.1 | 2.26 |
| Maximum predator approach speed | 6 | 972.9 | 3.10 |
| Maximum predator approach speed + Treatment | 7 | 975.2 | 5.37 |
| Maximum predator approach speed x Treatment | 8 | 977.5 | 7.64 |
| Baseline model (Reaction distance only) | 5 | 1002.7 | 32.90 |

**Table S5:** Results of Gamma GLMMs explaining variation in the maximum turning speed of the predator during the pursuit phase, based on 116 observations of 19 individual fish in trials where prey escaped at an acute angle (< 90°). Only the model including trial number represented a substantial improvement over the null model, and as trials progressed, fish reached higher maximum turning speeds. The analysis was based on 116 observations, not 117, as the maximum turning speed could not be calculated in one of the trials where the predator always moved a distance of less than 0.5 cm between successive video frames (heading angles were only calculated when the predator had moved a distance greater than 0.5 cm between frames).

| **Explanatory variables** | **Degrees of freedom** | **AICc** | **ΔAICc** |
| --- | --- | --- | --- |
| Trial number | 5 | 1418.0 | 0.00 |
| Treatment x Trial number | 7 | 1418.8 | 0.75 |
| Maximum predator approach speed x Treatment | 7 | 1425.6 | 7.55 |
| Maximum predator approach speed | 5 | 1425.7 | 7.64 |
| Maximum predator approach speed + Treatment | 6 | 1426.3 | 8.24 |
| Prey escape angle | 5 | 1427.2 | 9.20 |
| Null model (no explanatory variables) | 4 | 1427.3 | 9.27 |
| Treatment + Trial number | 6 | 1431.9 | 13.90 |
| Treatment | 5 | 1442.8 | 24.79 |

**Table S6:** Results of LMMs explaining variation in the minimum turn radius of the predator during the pursuit phase, based on 115 observations of 19 individual fish in trials where prey escaped at an acute angle (< 90°). The analysis was based on 115 observations, not 116, as the minimum turn speed could not be calculated in an additional trial where the prey was captured within two video frames (a minimum of three frames were required to calculate minimum turn radii).

| **Explanatory variables** | **Degrees of freedom** | **AICc** | **ΔAICc** |
| --- | --- | --- | --- |
| Prey escape angle | 4 | 1679.5 | 0.00 |
| Treatment + Prey escape angle | 5 | 1681.0 | 1.51 |
| Trial number | 4 | 1682.0 | 2.51 |
| Treatment x Prey escape angle | 6 | 1682.2 | 2.68 |
| Null model (no explanatory variables) | 3 | 1683.2 | 3.68 |
| Treatment + Trial number | 5 | 1684.0 | 4.53 |
| Treatment | 4 | 1685.1 | 5.59 |
| Maximum predator approach speed | 4 | 1685.3 | 5.81 |
| Treatment x Trial number | 6 | 1685.7 | 6.18 |
| Treatment + Maximum predator approach speed | 5 | 1687.3 | 7.76 |
| Treatment x Maximum predator approach speed | 6 | 1687.3 | 7.80 |
